# Neuronal hemoglobin induces loss of dopaminergic neurons in mouse Substantia nigra, cognitive deficits and cleavage of endogenous α-synuclein

**DOI:** 10.1101/2021.10.13.464204

**Authors:** Santulli Chiara, Bon Carlotta, De Cecco Elena, Codrich Marta, Narkiewicz Joanna, Parisse Pietro, Perissinotto Fabio, Santoro Claudio, Persichetti Francesca, Legname Giuseppe, Espinoza Stefano, Gustincich Stefano

## Abstract

**Backgroud:** Parkinson’s disease (PD) presents the selective loss of A9 dopaminergic (DA) neurons of Substantia Nigra *pars compacta* (SNpc) and the presence of intracellular aggregates called Lewy bodies. α-synuclein (α-syn) species truncated at the carboxy terminal (C-terminal) accumulate in pathological inclusions and promote α-syn aggregation and toxicity.

Hemoglobin (Hb) is the major oxygen carrier protein in erythrocytes. In addition, Hb is expressed in A9 DA neurons where it influences mitochondrial activity. Hb overexpression increases cells’ vulnerability in a neurochemical model of PD *in vitro* and forms cytoplasmic and nucleolar aggregates upon short-term overexpression in mouse SNpc.

**Methods:** α and β-globin chains were co-expressed in DA cells of SNpc *in vivo* upon stereotaxic injections of an Adeno-Associated Virus isotype 9 (AAV9) and in DA iMN9D cells *in vitro*.

**Results:** Long-term Hb over-expression in SNpc induced the loss of about 50% of DA neurons, a mild motor impairment and deficits in recognition and spatial working memory. Hb triggered the formation of endogenous α-synuclein C-terminal truncated species. Similar α-syn fragments were found *in vitro* in DA iMN9D cells over-expressing α and β-globins when treated with pre-formed α-syn fibrils.

**Conclusion:** Our study positions Hb as a relevant player in PD pathogenesis for its ability to trigger DA cells’ loss *in vivo* and the formation of C-terminal α-synuclein fragments.

## BACKGROUND

Parkinson’s disease (PD) is a chronic progressive neurodegenerative disorder clinically defined in terms of motor symptoms. The most evident pathological hallmarks are the selective degeneration of A9 dopaminergic (DA) neurons of Substantia Nigra *pars compacta* (SNpc) and the presence of intracellular aggregates called Lewy bodies (LB). The consequent loss of DA synapses in the striatum is the primary origin of the inability to control movements [1]. The causes of the selective degeneration of A9 neurons remain largely unknown.

Highly penetrant mutations producing rare, monogenic forms of the disease are in genes involved in mitochondria homeostasis, vesicular trafficking and lysosomal function [2]. The discovery of missense mutations or duplication/triplication of the α-syn gene (SCNA) proves its causative role in autosomal-dominant early-onset PD [3]. α-syn protein is the major constituent of LB with a mix of the full-length protein (FL-α-syn) and of C-terminal truncated fragments (ΔC-α-syn). Progressive α-synuclein aggregation can be recapitulated in model systems by exogenous introduction of *in vitro* generated preformed fibrils (PFFs) [4–6]. PFFs seed the aggregation of endogenous α-synuclein with a prion-like mechanism and this aggregation may spread along synaptically connected pathways [4, 7].

In the quest of the molecular basis of neurodegeneration, the vulnerability of DA cells of the SNpc is associated to genes differentially expressed in comparison to DA neurons of the Ventral Tegmental Area (VTA; A10), largely spared in disease. Transcripts and encoded proteins for α-and β-chains of Hemoglobin (Hb) are enriched in A9 DA neurons of mouse SNpc [8–10]. This pattern of expression is conserved in the human mesencephalic DA cells’ system [11]. Neuronal Hb retains its α2β2 tetrameric structure as in blood where it exerts its function as oxygen-carrier molecule [10]. In human brain, α-and β-chains are co-localised in the mitochondrion and interact with mitochondrial proteins [12]. Mitochondrial Hb is significantly reduced in aged monkey striatum and human SN [13, 14] as well as in PD post-mortem brains where it may form complexes with α-syn [14, 15].

The co-overexpression of α-and β-chains in the mouse DA iMND9 cell line *in vitro* and in mouse SNpc by AAV-mediated delivery *in vivo* has proved that Hb may interfere with cellular pathways targeted in PD. *In vitro* Hb overexpression alters transcript levels of genes involved in oxygen (O_2_) homeostasis and oxidative phosphorylation suggesting a role in mitochondrial activity [8]. It also increases DA cells’ susceptibility to 1-methyl-4-phenylpyridinium (MPP_+_) and rotenone [9], neurochemical *in vitro* models of PD targeting mitochondrial complex I activity [16]. Upon stress, Hb inhibits autophagy as well as it moves to the nucleus where it forms insoluble aggregates in the nucleolus triggering nucleolar stress [9].

When AAV carrying α- and β-chains of Hb are stereotaxically injected into mouse SN, after one-month Hb forms cytoplasmic and nucleolar aggregates [9]. While no effects on DA neurons number are evident, a motor learning impairment is observed, urging the need for further analysis at longer time points.

Here, we show that a prolonged ectopic expression of Hb in rodent SNpc induces the loss of DA neurons along with motor and cognitive impairments. Linking Hb to molecular pathways involved in PD, Hb overexpression *in vivo* and *in vitro* provokes the formation of ΔC-α-syn, a post-translational modification associated to PD. These results provide new evidence in support of the role of Hb in PD pathogenesis.

## MATERIAL AND METHODS

### Animals

All animal experiments were performed in accordance with European guidelines for animal care and following Italian Board Health permissions (D.Lgs. 26/2014, 4 March 2014). Mice were housed and bred in IIT – Istituto Italiano della Tecnologia (Genova, GE, Italy) animal facility, with 12 hours dark/night cycles and controlled temperature and humidity. Food and water were provided *ad libitum*.

### Behavioural testing

All procedures involving animals and their care were carried out in accordance with the guidelines established by the European Community Council (Directive 2010/63/EU of September 22, 2010) and were approved by the Italian Ministry of Health (DL 116/92; DL 111/94-B).

### Locomotor activity

To measure spontaneous locomotor activity, mice were placed in the locomotor activity chambers (Omnitech Digiscan, Accuscan Instruments, Columbus, ОН, USA) for 60 minutes and total distance travelled was measured by the analysis of infrared beam interruptions.

### Rotarod

A Rotarod from TSE Systems was used. Briefly, mice were handled on alternate days during the week preceding the start of the Rotarod test (3 handling sessions; 1 min per mouse per session). Behavioural testing lasted two days. On day 1, mice habituation to rotation on the rod under a constant speed of 4 rpm for three trials (60-s inter-trial interval) were performed in the morning. Trial ended as the mouse was falling off the rod or if it was spending on the rod more than 300 s. In the afternoon, mice were placed on the rod having a constant 4 rpm-speed for 60 seconds. Then, the accelerating program was launched for three trials (60-s inter-trial interval). Trial ended as previously described. Mice were tested the day after only as for Day1 afternoon. Time spent on the rod was automatically recorded. The average time spent on the rod was then calculated.

### Static rods

Five 60 cm long wooden rods of varying thickness (35, 25, 15, 10 and 8 mm diameter) were fixed to a laboratory shelf horizontally protruding into space at 60 cm of height above the floor. Mice were placed outward at the far end of the widest rod. Two measures were considered: orientation time (time taken to orientate 180° from the starting position towards the shelf) and transit time (the time taken to travel to the shelf end). Orientation was dependent on the mouse staying upright. The maximum score of 120 sec was assigned when mouse turned upside down and clung below the rod, fell or reached the maximum test time. If the mouse fell off the rod within 5 seconds, another attempt was allowed (as falling within 5 sec could be due to faulty placing by the experimenter), for a maximum of three trials, and the best result was considered. Mice were not tested on smaller rods. Once tested on one rod, mice were placed back to the home cage to rest, while other mice were tested. This procedure was repeated for all the rods. The time for orientation and transit were plotted in the graphs for statistical analysis. Since in smaller rods many of the mice fell or did not complete the test, the success rate of the test was also calculated as the number of mice that completed the test and the Chi-square test was used to compare the two groups of mice.

### Horizontal bars

For this test, two bars made of brass were used, 40 cm long, held 50 cm above the bench surface by a wooden support column at each end. Two bar diameters were used: 2 and 4mm. As mice find it easiest to grasp the narrow 2 mm bar, mice were first tested on this bar. Holding it by the tail, the mouse was placed on the bench in front of the apparatus, pulled quickly backwards about 20 cm (perpendicular alignment to the bar), rapidly raised and allowed to grasp the horizontal bar at the central point with its forepaws only and then the tail was released. The time to reach one of the edges of the bar was recorded. If the mouse had failed to grasp the bar properly at the first attempt or fell within 5 sec, the score was not recorded, and it was placed back to the cage to rest and then the trial was repeated up to three times since a poor placement of the operator might occur. The best score was excluded from these additional trials. The score was calculated as an average of the scores of the 2mm and 4 mm bar trials as indicated below:

Falling between 1-5 sec = 1

Falling between 6-10 sec = 2

Falling between 11-20 sec = 3

Falling between 21-30 sec = 4

Falling after 30 sec = 5

The maximum score for completing the test was 5 for each bar and 10 for both bars.

### Novel object recognition test

Mice were handled on alternate days during the week preceding the start of the test (3 handling sessions; 1 min per mouse per session on Day 1, 3, 5.). On day 6, mice were subjected to the habituation session in the empty open field for 1 hour. The intensity of the light on the apparatus was of about 60 lux. On day 7, each mouse was subjected to two successive sessions. Pre-test session (acquisition trial, 10 minutes): each mouse was introduced into the open field containing two identical copies of the same object. At the end of the session the mouse was placed into the home cage. Test session (retention trial, 5 minutes): 1 hour after the acquisition trial, both objects were substituted with a third copy of previous object and a new object. The animals were considered to be exploring the object when they were facing (at a distance < 1 cm) or were touching or sniffing the object. Mice that explored object for <4 sec were excluded. The type and positions of presentation of the objects during acquisition and retention phase were counterbalanced across animals. The preference index, expressed as the ratio of the amount of time spent exploring the objects (training session) or the novel one (retention session) over the total time spent exploring both objects, defined the recognition memory. In the pre-test session, the total amount of exploration (in sec) for the identical objects was used to verify no preference for one of the two side of the chamber where the objects were located.

### Y-maze Spontaneous Alternation Test

The apparatus was a Y-shaped maze with three opaque arms spaced 120° apart with a measure of 40 × 8 × 15cm each. An overhead camera was mounted to ceiling directly above apparatus to monitor mice movement and 4 standing lamps with white light bulbs were placed at the corners outside privacy blinds pointed away from apparatus.

The arms were labelled as A, B or C to identify the entries. The animal was placed inside arm B facing away from centre and allowed to move through apparatus for 10 minutes while being monitored by automated tracking system. Trial began immediately and ended when defined duration had elapsed. Scoring consisted of recording each arm entry (defined as all four paws entering arm). The total entries in all arms were recorded. A spontaneous alternation occured when a mouse entered a different arm of the maze in each of 3 consecutive arm entries. The % of spontaneous alternation was then calculated as ((#spontaneous alternation/(total number of arm entries-2))×100.

### Stereotaxic AAV9 injection

Adult (12 weeks old) male C57BI/6J mice were used for experiments. Mice were anesthetized by a mixture of Isoflurane/Oxygen and placed on a stereotaxic apparatus (David Kopf instrument, Tujunga, CA, USA) with mouse adaptor and lateral ear bars. The skin on the skull was cut and one hole was made on both side by a surgical drill. A stereotaxic injection of 1 μl of viral vector suspension (AAV9-CTRL or a mixture of AAV9-2xFLAG-α-globin and AAV9-β-globin-MYC, called AAV9-Hb; titer: – 5*10^12^ vg/ml) was delivered bilaterally to SNpc at the following coordinates: anterior/posterior (A/P) −3.2 mm from bregma, medio/lateral (M/L) −1.2 mm from bregma and dorso/ventral (D/V) − 4.5 mm from the dura. The coordinates were calculated according to the Franklin and Paxinnos Stereotaxic Mouse Atlats. Injection rate was 1 μl/15 minutes using a glass gauge needle. After the infusion, the needle was maintained for another 1 minute in the same position and then retracted slowly.

### Tissue collection and processing

At 10 months after injection of AAVs into SNpc, the animals were sacrificed. Following induction of deep anaesthesia with an overdose of a mixture of Xylazina and Zoletil, the animals were intensively perfused transcardially with PBS 1 ×. For biochemical analysis, SNpc was dissected and immediately frozen in liquid nitrogen and stored at −80 °C, pending analyses. For immunohistochemical analysis, after the intensively transcardially perfusion with PBS 1X, animals were perfused with 4% paraformaldehyde diluted in PBS 1X. Brains were postfixed in 4% paraformaldehyde for 1 h at 4 °C. The regions containing the SN were cut in 40 μm free-floating slides with a vibratome (Vibratome Series 1000 Sectioning System, Technical Products International, St. Louis, MO, USA). Four consecutive series were collected to represent the whole area of interest.

### Immunofluorescence with labelled α-syn fibrils

Human α-syn fibrils were labelled with Alexa-488 succinimidyl esther (Thermo Fisher Scientific, A20000) following manufacturer’s instructions and the unbound fluorophore was removed with multiple dialysis steps in sterile PBS. Uptake experiments were performed following standard IF protocol or following the protocol described by Karpowicz *et al.* [35]. Briefly, cells seeded on coverslips were incubated with culture medium containing labelled α-syn fibrils for 24 h. Prior to standard immunocytochemistry protocol, fluorescence from non-internalized fibrils was quenched by incubating with Trypan Blue for 5 minutes. Cells were then fixed in 4% paraformaldehyde for 20 minutes, washed two times and permeabilized with 0.1% Triton X-100 in PBS 1X for 4 minutes and incubated with HCS Blue Cell Mask 1:1000 for 30 minutes (Thermo Fisher Scientific). Cells were washed twice in PBS 1X and once in Milli-Q water and mounted with Vectashield mounting medium (Vector Lab, H-1000). Images acquisition was performed using C1 Nikon confocal microscope (60x oil, NA 1.49, 7x zoom-in) as z-stacks of 0.5 μm.

### Immunocytochemistry

Cells were washed twice with D-PBS, fixed in 4% paraformaldehyde for 20 minutes, washed twice with PBS 1X and treated with 0.1M glycine for 4 minutes in PBS 1X, washed twiceand permeabilized with 0.1% Triton X-100 in PBS 1X for 4 minutes. Cells were then incubated in blocking solution (0.2% BSA, 1% NGS, 0.1% Triton X-100 in PBS 1X), followed by incubation with primary antibodies diluted in blocking solution for 2:30 hours at room temperature. After two washes in PBS 1X, cells were incubated with labelled secondary antibodies and 1μg/ml DAPI (for nuclear staining) for 60 minutes. Cells were washed twice in PBS 1X and once in Milli-Q water and mounted with Vectashield mounting medium (Vector Lab, H-1000). The following antibodies were used: anti-FLAG 1:100 (Sigma–Aldrich, F7425), anti-MYC 1:250 (Cell Signaling, 2276), anti-α-syn (C-20) 1:200 (Santa Cruz Biotechnology, sc-7011-R), anti-α-syn (SYN-1) 1:200 (BD Transduction Laboratories, 610787) and anti-α-syn(phosphoS129) 1:200 (Abcam, ab59264). For detection, Alexa Fluor-488, −594 or −647 (Life Technologies) antibodies were used. Image acquisition was performed using C1 Nikon confocal microscope (60x oil, NA 1.49, 7x zoom-in).

### Immunohistochemistry

For immunohistochemistry, free-floating slides were rinsed three times in 0.1M phosphate buffered saline (PBS; pH 7.6), contained 0.1% Triton X-100 between each incubation period. All sections were quenched with 3% H_2_O_2_/10% for 10 min, followed by several changes of buffer. As a blocking step, sections were then incubated in 7% normal goat serum and 0.1% Triton-X 100 for 2 hours at room temperature. This was followed by incubation in primary antibody diluted in 3% normal goat serum and 0.1% Triton-X 100 at 4°C for 24 hrs. The antibody used was an anti-TH diluted 1:500 (AB-152, Millipore). After over-night incubation with the primary antibody, sections were rinsed and then incubated for 2 hours at room temperature with biotinylated secondary antibodies (anti-rabbit 1:1000; Thermo Scientific) in the same buffer solution. The reaction was visualized with avidin-biotin-peroxidase complex (ABC-Elite, Vector Laboratories), using 3,3-diaminobenzidine as a chromogen. Sections were mounted on super-frost ultra plus slides (Thermo Scientific), dehydrated in ascending alcohol concentrations, cleared in xylene and coverslipped in DPX mounting medium.

For fluorescent immunohistochemistry, free-floating slides were treated with 0.1 M glycine for 5 min in PBS 1 × and then with 1% SDS in PBS 1 × for 1 min at RT. Slides were blocked with 10% NGS, 1% BSA in PBS 1 × for 1 h at RT. The antibodies were diluted in 1% BSA, 0.3% Triton X-100 in PBS 1 ×. For double immunoflurescence, incubation with primary antibodies was performed overnight at RT and incubation with 1:500 Alexa fluor-conjugated secondary antibodies (Life Technologie) was performed for 2 h at RT. Nuclei were labelled with 1 μg/ml DAPI. Slides were mounted with mounting medium for fluorescence Vectashield (Vector Laboratories). The following primary antibodies were used: anti-TH 1:1000 (Sigma-Aldrich or Millipore), anti-FLAG 1:100 (Sigma-Aldrich), anti-MYC 1:100 (Cell Signaling) and anti-Hemoglobin 1:1000 (MP Biomedicals). For detection, Alexa fluor-488 or −594 (Life Technologies) were used. All images were collected using confocal microscopes (LEICA TCS SP2).

### Quantification of DA neurons in the SNPc

The number of TH positive cells were determined by counting every fourth 40-μm sections as previously described [17]. The delimitation between the ventral tegmental area and the SN was determined by using the medial terminal nucleus of the accessory optic tract as a landmark. All counts were performed blind to the experimental status of the animals through ImageJ software. TH_+_ cells were counted using “3D object counter tool”. Each found object has been quantified applying default settings. The following parameters were modified: Size filter set to 10-20 voxels, threshold set to: 128. Values were expressed as absolute quantification of unilateral SNpc TH_+_ cells.

### Statistical Analysis

All data were obtained by at least three independent experiments. Data represent the mean ± S.E.M. and each group was compared individually with the reference control group using GraphPad Prism (v9) software. To compare the means of two samples, groups were first tested for normality, and then for homogeneity of variance (homoscedasticity). If the normality assumption was not met, data were analysed by nonparametric Mann-Whitney test. If the normality assumption was met, but homogeneity of variance was not, data were analysed by unpaired two-tailed t-test followed by Welch’s correction. If both assumptions were met, data were analysed by unpaired two-tailed t-test. To compare more than 2 groups One-Way ANOVA was used. Statistical analysis of static rods experiments, each group were analysed by Chi-squared test. Significance to reference samples are shown as *, p ≤ 0.05; **, p ≤ 0.01; ^***^, p≤ 0.001; ****, p ≤ 0.0001.

### Western Blot

iMN9D cells were washed 2 times with D-PBS and lysed in 300 μl SDS sample buffer 2X (6 well-plate), briefly sonicated, boiled and 10 μl/sample loaded on 15 % or 8 % (for Spectrin α II immunoblot) SDS-PAGE gel. For antibodies validation, cells were lysed in cold lysis buffer (10 mM Tris-HCl pH 8, 150 mM NaCl, 0.5% Igepal CA-630, 0.5% sodium deoxycholate) supplemented with protease inhibitor mixture (Roche Diagnostics, COEDTAF-RO). Lysates were incubated for 30 minutes at 4 °C on rotator and cleared at 12000xg for 20 minutes at 4 °C. Supernatants were transferred in new tubes and total protein content was measured using bicinchoninic acid protein (BCA) quantification kit (Pierce) following the manufacturer’s instructions. For SNpc lysates, dissected brain area was lysed in cold RIPA buffer and centrifuged for 10 minutes at 17,000xg. Sample buffer was added to the supernatant and boiled at 95°C for 5 minutes and 30 μg of proteins were loaded on a 10% SDS-PAGE gel. Proteins were transferred to nitrocellulose membrane (Amersham™, Cat. No. GEH10600001) for 1:30 hour at 100V or 16 hours 20V (only for Spectrin α II immunoblot). Membranes were blocked with 5% non-fat milk or 5% BSA (only for Spectrin ⍺ II immunoblot) in TBST solution (TBS and 0.1% Tween20) for 40 minutes at room temperature. Membranes were then incubated with primary antibodies at room temperature for 2 h or overnight at 4 °C (only for Spectrin α II immunoblot). The following antibodies were used: anti-FLAG 1:2000 (Sigma–Aldrich, F3165), anti-MYC 1:2000 (Cell Signaling, 2276), anti-β-actin 1:5000 (Sigma–Aldrich, A5441), anti-Hemoglobin 1:1000 (Cappel, MP Biomedicals, 55039) and anti-GFP 1:1000 (Clontech, 632380), anti-α-syn 1:1000 (C-20) (Santa Cruz Biotechnology, sc-7011-R), anti-α-syn 1:1000 (SYN-1) (BD Transduction Laboratories, 610787), anti-biotin-HRP (Jackson ImmunoResearch Laboratories), anti-Spectrin ⍺ II 1:1000 (Santa Cruz, Cat. No. sc-46696), anti-TH 1:1000 (Millipore). For development, membranes were incubated with secondary antibodies conjugated with horseradish peroxidase (Dako) for 1 hour at room temperature. For IP and pulldown experiments, membranes were incubated with Protein A antibody conjugated with horseradish peroxidase for 1 hour at room temperature. Proteins of interest were visualized with the Amersham ECL Detection Reagents (GE Healthcare by SIGMA, Cat. No. RPN2105) or LiteAblot TURBO Extra-Sensitive Chemioluminescent Substrate (EuroClone, Cat. No. EMP012001). Western blotting images were acquired using with Alliance LD2-77WL system (Uvitec, Cambridge) and band intensity was measured UVI-1D software (Uvitec, Cambridge).

### Production of recombinant human α-syn

Expression and purification of human α-syn were performed as previously described [18]. Briefly, α-syn cDNA was cloned in pET-11a vector and expressed in *E.coli* BL21(DE3) strain. Cells were grown in Luria-Bertani medium at 37°C and expression of α-syn was induced by addition of 0.6 mM isopropyl-β-D-thiogalactoside (IPTG) followed by incubation at 37°C for 5 hours. The protein was extracted and purified according to Huang *et al*. [19].

### Fibrillation of human α-syn

Lyophilized human α-syn was re-suspended in ddH_2_O, filtered with a 0,22 μm syringe filter and the concentration was determined by absorbance measured at 280 nm. Fibrillization reactions were carried out in a 96-well plate (Perkin Elmer) in the presence of a glass bead (3 mm diameter, Sigma) in a final reaction volume of 200 μL. Human α-syn (1.5 mg/ml) was incubated in the presence of 100 mM NaCl, 20 mM TrisHCl pH 7.4 and 10 uM thioflavin T (ThT). Plates were sealed and incubated in BMG FLUOstar Omega plat reader at 37°C with cycles of 50 seconds shaking (400 rpm) and 10 seconds rest. Formation of fibrils was monitored by measuring the fluorescence of ThT (excitation: 450 nm, emission: 480 nm) every 15 minutes. After reaching the plateau phase, the reactions were stopped. Fibrils were collected, centrifuged at 100000g for 1 hour, resuspended in sterile PBS and stored at −80°C for further use. For cell culture experiments, the fibrillation reaction was carried out without ThT and PFFs in 0,5 ml conical plastic tubes were sonicated for 5 minutes in a Branson 2510 Ultrasonic Cleaner prior addition to cell culture medium.

### Cleaning procedures

For fibrils inactivation, all contaminated surfaces and laboratory wares, both reusable and disposable, were cleaned using a 1 % SDS solution prior washing with Milli-Q water according to Bousset *et al.* [20].

### Atomic Force Microscopy

Atomic Force Microscopy (AFM) was performed as previously described [21]. Briefly, three to five μl of fibril solution was deposited onto a freshly cleaved mica surface and left to adhere for 30 minutes. Samples were then washed with distilled water and blow-dried under a flow of nitrogen. Images were collected at a line scan rate of 0.5-2 Hz in ambient conditions. The AFM free oscillation amplitudes ranged from 25 nm to 40 nm, with characteristic set points ranging from 75% to 90% of these free oscillation amplitudes.

### Cell line

MN9D-Nurr1^Tet-on^ (iMN9D) cell line stably transfected with pBUD-IRES-eGFP (CTRL cells) or with pBUD-β-globin-MYC IRES-eGFP, 2xFLAG-α-globin (Hb cells) were used [8]. Cells were maintained in culture at 37 °C in a humidified CO2 incubator with DMEM/F12 medium (Gibco by Life Technologies, DMEM GlutaMAX® Supplement Cat. No. 31966-021; F-12 Nutrient Mix GlutaMAX® Supplement Cat. No. 31765-027) supplemented with 10% fetal bovine serum (Euroclone, Cat. No. ECS0180L), 100 μg/ml penicillin (Sigma–Aldrich), 100 μg/ml streptomycin (Sigma-Aldrich). 300 μg/ml neomycin (Gibco by Life Technologies, Cat. No 11811-031) and 150 μg/ml zeocyn (Invivogen, Cat. No. ant-zn-05) were used for selection.

### Exposure of iMN9D cells to α-syn monomers and fibrils

iMN9D cells were exposed to 2 μM of α-syn species (2 μM equivalent monomer concentration in the case of amyloids) in cell culture media for 24, 48, 96 hours before collection.

For western blot analysis cells were plated in 6 well-plate (6×10^5^ cells/plate for 24h collection, 3×10^5^ cells/plate for 48h collection, 2×10^5^ cells/plate for 96h collection). Additionally, at 96 hours cells were split and maintained for three additional days before collection as Passage 1 (P1). Cells treated with vehicle were used as control (Untreated).

For immunocytochemistry, cells were cultured in 12-well plates with coverslips (3×10^5^ cells/well for 24h collection, 1,5×10^5^ cells/well for 48h collection, 1×10^5^ cells/well for 96h collection).

### Pull-down assay of biotinylated fibrils

For pulldown experiments, α-syn PFFs were biotinylated following the manufacturer’s instructions (Sigma-Aldrich). iMN9D cells were lysed in cold immunoprecipitation (IP) buffer (50 mM Tris HCl pH 8, 150 mM NaCl, 0.1% Igepal CA-630) containing protease inhibitors (Roche Diagnostics, COEDTAF-RO). Following 30 min incubation at 4 °C on rotator, lysates were cleared at 12000 g for 20 min and incubated with biotinylated fibrils overnight at 4°C on rotator. Biotinylated fibrils were pulled down by binding to NeutrAvidin Agarose Resin (Pierce, 29200). After 4 h incubation at 4 °C, the resin-bound complexes were washed three times with IP buffer and eluted with SDS sample buffer 2X, boiled at 95°C for 5 min and analysed by western blot.

### Cathepsin D and Calpain inhibitors treatment

Calpain inhibitor III (Santa Cruz Cat. No. SC-201301) and Pepstatin A (Pep, Cathepsin D inhibitor, MedChem Express Cat. No. HY-P0018) were dissolved in dimethylsulfoxide (DMSO) and diluted in cell culture medium to a final concentration respectively of 10 μM and 100 μM. Hb cells were treated with vehicle (DMSO), Calpain inhibitor III and Pepstatin A at the indicated concentrations for 24 h. Medium was then removed and replaced with new one containing α-syn PFFs, as previously reported, and protease inhibitors to a final concentration respectively of 10 μM and 100 μM, as the day before. Cells were collected at the indicated time points for western blot analysis. Immunoblot of Spectrin ⍺ II was used to monitor Calpain inhibitor activity, while Cathepsin D activity kit was used to monitor Cathepsin D activity.

### Cathepsin D activity assay

Cathepsin D (CatD) activity measurements were performed using the Cathepsin D activity assay kit (BioVision, Cat. N. K143) following manufacturer’s instructions. Briefly, cells were washed twice with PBS, collected in culture media and pelleted by centrifugation at 500 g for 5 min. Cells were counted and 1×10 ^5^ cells/well were used. Cells were washed once with PBS, pelleted again by centrifugation at 500 g for 5 min and lysed in CD Cell Lysis Buffer incubating samples for 10 min on ice. Cells were then centrifuged at maximum speed for 10 min. As control, untreated cells were incubated with PepA (100 μM final concentration) at 37°C for 10 min prior addition of Reaction Buffer and CD substrate (Positive control). The reaction was left to proceed at 37°C for 1:30 h in the dark. Fluorescence was read using Thermo Scientific Varioskan® Flash with a 328-nm excitation filter and 460-nm emission filter. CD activity in relative fluorescence units (RFU) was then normalized to Hb cells treated with vehicle and indicated as % Activity. Each sample was measured in duplicate and measurements were repeated two times.

### RNA isolation, Reverse Transcription (RT) and quantitative RT-PCR (qRT-PCR)

Total RNA was extracted using TRIzol Reagent (Thermo Fisher, 15596026) and following manufacturer’s instructions. RNA samples were subjected to TURBO DNase (Invitrogen, Cat. No. AM1907) treatment, to avoid DNA contamination. The final quality of RNA sample was tested on 1 % agarose gel with formaldehyde. A total of 1 μg of RNA was subjected to retrotranscription using iScript™cDNA Synthesis Kit (Bio-Rad, Cat. No. 1708890), according to manufacturer’s instructions. qRT-PCR was carried out using SYBR green fluorescent dye (iQ SYBR Green Super Mix, Bio-Rad, Cat. No. 1708884) and an iCycler IQ Real time PCR System (Bio-Rad). The reactions were performed on diluted cDNA (1:2). Mouse actin was used as normalizing control in all qRT-PCR experiments. The amplified transcripts were quantified using the comparative Ct method and the differences in gene expression were presented as normalized fold expression with ΔΔCt method [36, 37]. The following primer pairs were used:

β-actin: fwd CACACCCGCCACCAGTTC, rev CCCATTCCCACCATCACACC;

Capn1: fwd TTGACCTGGACAAGTCTGGC, rev CCGAGTAGCGGGTGATTATG;

Capn2: fwd ATGCGGAAAGCACTGGAAG, rev GACCAAACACCGCACAAAAT;

Ctsd: fwd CAGGACACTGTATCGGTTCCA, rev CAAAGACCGGAAGCACGTTG.

## RESULTS

### Long-term Hb overexpression in SNpc triggers loss of DA neurons, decreases motor performances and causes cognitive impairments

To study the effects of Hb overexpression on endogenous α-syn *in vivo*, we injected a mixture of AAV9-2xFLAG-α-globin and AAV9-β-globin-MYC (indicated as AAV9-Hb) or AAV9-CTRL bilaterally into the SNpc of 3 months old mice (respectively named as Hb and CTRL mice). In a previous study [9], over-expression of Hb for one month in SNpc of mice infected with the very same AAV9-Hb provoked the formation of cytoplasmic and nucleolar Hb aggregates and a mild motor learning impairment preserving DA cells viability. To investigate long-term Hb effects on DA neurons’ homeostasis and mice behaviour, here we prolonged Hb overexpression up to 9 months after the injection (Figure 1a).

**Figure 1.**
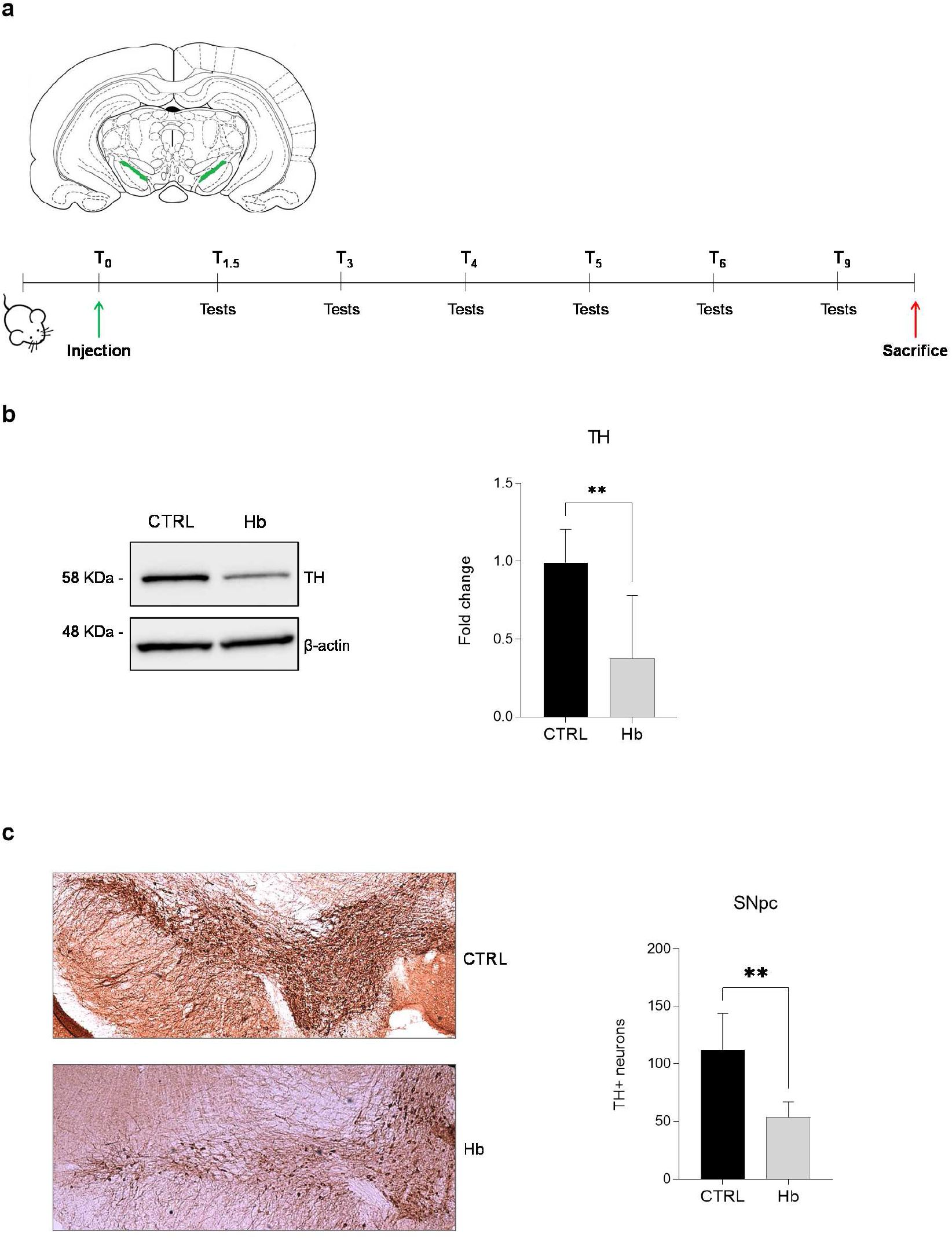
Hb overexpression in SNpc triggers the loss of DA neurons. Scheme representing the experimental protocol used for the assessment Hb overexpression. AAV9 expressing Hb (Hb) and CTRL (CTRL) constructs were bilaterally injected in the brain of 3-months old mice. Brain diagram indicating SNpc (green) as the region of the injection (*upper panel*). Behavioural tests were performed to verify the locomotor performance of mice 1.5, 3, 4, 5, 6 and 9 months post-injection (*lower panel*) (**a**). Level of expression in SNpc of Hb and CTRL mice (n=4) at 10 months post-injection of tyrosine hydroxylase (TH, 58 KDa) (**b**, *left panel*) was assessed by western blot. Band intensity was quantified (**b**, *right panel*). TH-positive neurons of the SNpc were evaluated by immunohistochemistry (**c**, *left panel*) and quantified (**c**, *right panel*) for Hb and CTRL mice (n=4; 3 slices each). Data represent means ± SEM. Statistical analysis was performed with unpaired t test with Welch’s correction. *, p ≤ 0.05; **, p ≤ 0.01; ^***^, p≤ 0.001; ****, p ≤ 0.0001.

Firstly, Hb mice presented a decrease of tyrosine hydroxylase (TH) expression of about 50% by western blot analysis (Figure 1b). To understand whether the decrease in TH levels was due to a loss of neurons or a change in expression, we performed immunohistochemistry analysis of brain tissues from Hb and CTRL mice and quantified DA cells in SNpc. A significant loss of about 50% of DA neurons was evident in SNpc of Hb mice proving that long-term overexpression of Hb was detrimental to well-being of DA neurons (Figure 1c).

To determine the behavioural impact of Hb overexpression, we monitored Hb and CTRL mice by a series of tests over a period of 9 months after the injection (Figure 1a). No gross difference in locomotor activity was observed between the two groups at all the time points examined (Figure 2a and b). However, specific deficits in Hb mice were evident by applying more subtle tests. In horizontal bars, a test that measures the forelimb strength and coordination, Hb mice were performing worse than CTRL animals starting from 5 months after the injection (Figure 2c). In static rods, a test to evaluate the coordination of mice to walk on wooden rods of different diameter, Hb mice performed worse on narrow rods (10 mm). Hb mice often fall out of the rods both during orientation and transit, and they failed to complete the test significantly more often than CTRL mice, in particular those at the 9 months post-injection time point (Figure d; Supplementary Figure 1).

**Figure 2.**
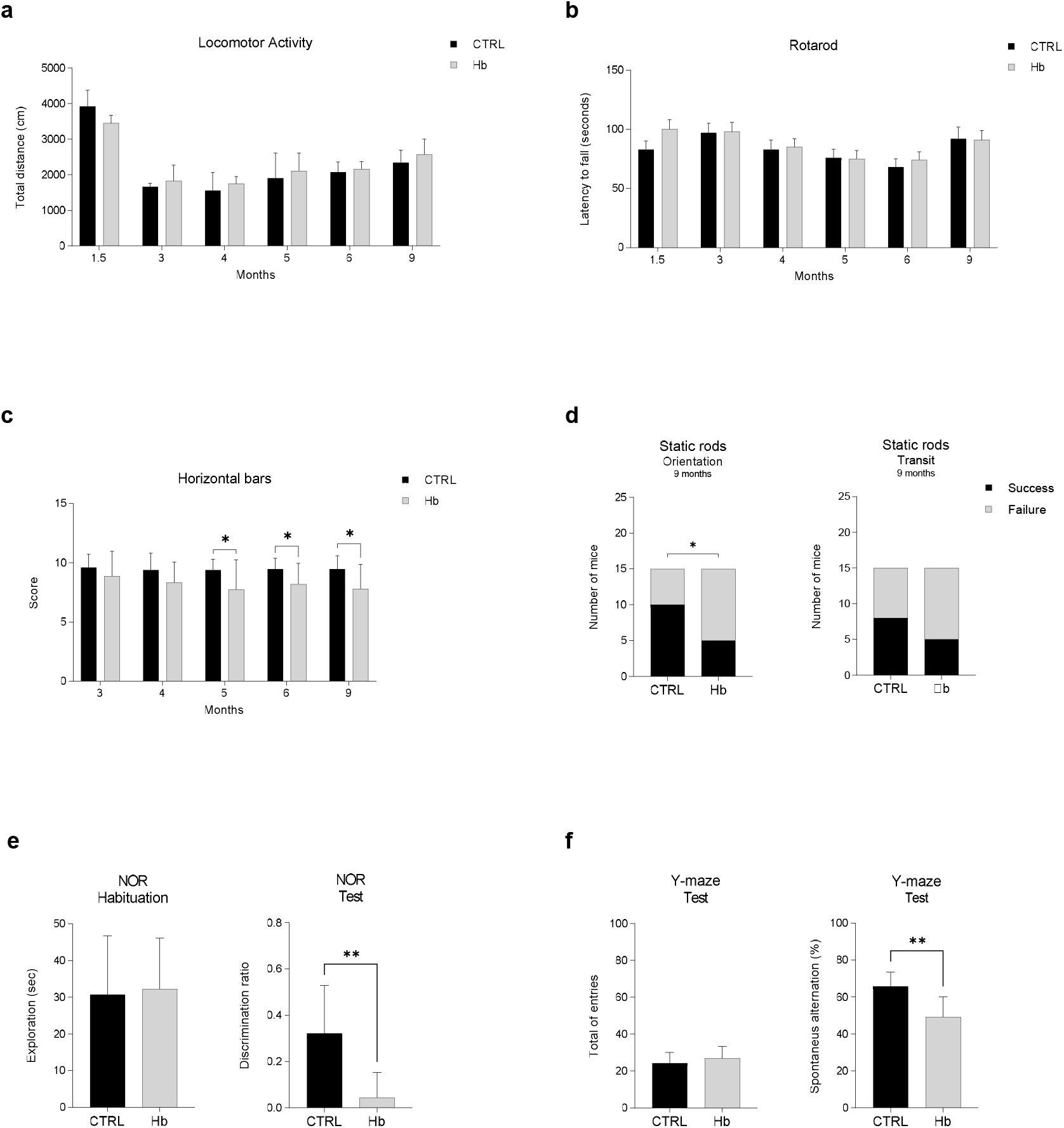
Hb overexpression in SNpc decrease motor performances and trigger cognitive impairments. CTRL (n=12) and Hb (n=12) mice were assessed for locomotor activity and total distance (cm) was recorded at different time points after injections (**a**). CTRL (n=15) and Hb (n=15) mice were also scored for motor coordination with the rotarod test (**b**) and latency to fall was measured. Horizontal bars test was used to assess forelimb strength and coordination and CTRL (n=15) and Hb (n=15) mice were scored for their performance(**c**). Data represent means ± SEM. Statistical analysis was performed with unpaired t test with Welch’s correction. *, p ≤ 0.05. CTRL (n=15) and Hb (n=15) were assessed in static rods test measuring two parameters, transit time and orientation time (seconds) and different time points. In panel (**d**) it is depicted the results at 9 months after injection. Chi-square test was used to evaluate the success/failure of each group. *, p ≤ 0.05. Novel object recognition test (NOR) was used to evaluate recognition memory (**e**). CTRL (n=8) and Hb (n=8) was habituated to the object for 10 minutes and exploration (seconds) of the objects was measured (**e**, *left panel*). After 1 hours, mice were assessed to recognize the novel object and the discrimination ration was plotted (**e**, *right panel*). Spontaneous alternation in the Y-maze was used to measured spatial working memory in CTRL (n=10) and Hb (n=12) mice (**f**). Total entries were calculated for each group (**f**, *left panel*). Spontaneous alternation % was plotted for each group (**f**, *right panel*). Data represent means ± SEM. Statistical analysis was performed with unpaired t test with Welch’s correction. *, p ≤ 0.05; **, p ≤ 0.01.

Since PD patients may experience cognitive dysfunctions [1], we tested mice in two cognitive tests, the novel object recognition test (NOR) and the Y-maze for spontaneous alternation, assessing recognition memory and spatial working memory, respectively. In the NOR, Hb mice showed a strong deficit in recognizing the novel object, as indicated by the discrimination ratio (Figure 2e). In the Y-maze, Hb mice displayed less spontaneous alternation compared to CTRL mice with no difference in the total entries in the arms (Figure 2f).

These data demonstrate that Hb expression in SNpc caused partial loss of DA neurons and induced mild motor impairments and significant cognitive deficits.

### Biochemical analysis of endogenous α-syn upon Hb overexpression in vivo

To characterize the effects of Hb overexpression in DA neurons of SNpc, we studied the metabolism of endogenous α-syn carrying out a western blot analysis with two epitope-specific antibodies: C-20 and SYN-1, recognising the full-length (FL-α-syn) or the C-terminal truncated (ΔC-α-syn) isoforms, respectively (Figure 3a). Interestingly, a statistically significant increase in ΔC-α-syn/FL-α-syn ratio was observed in the SNpc of mice overexpressing Hb compared to the control group (Figure 3b).

**Figure 3.**
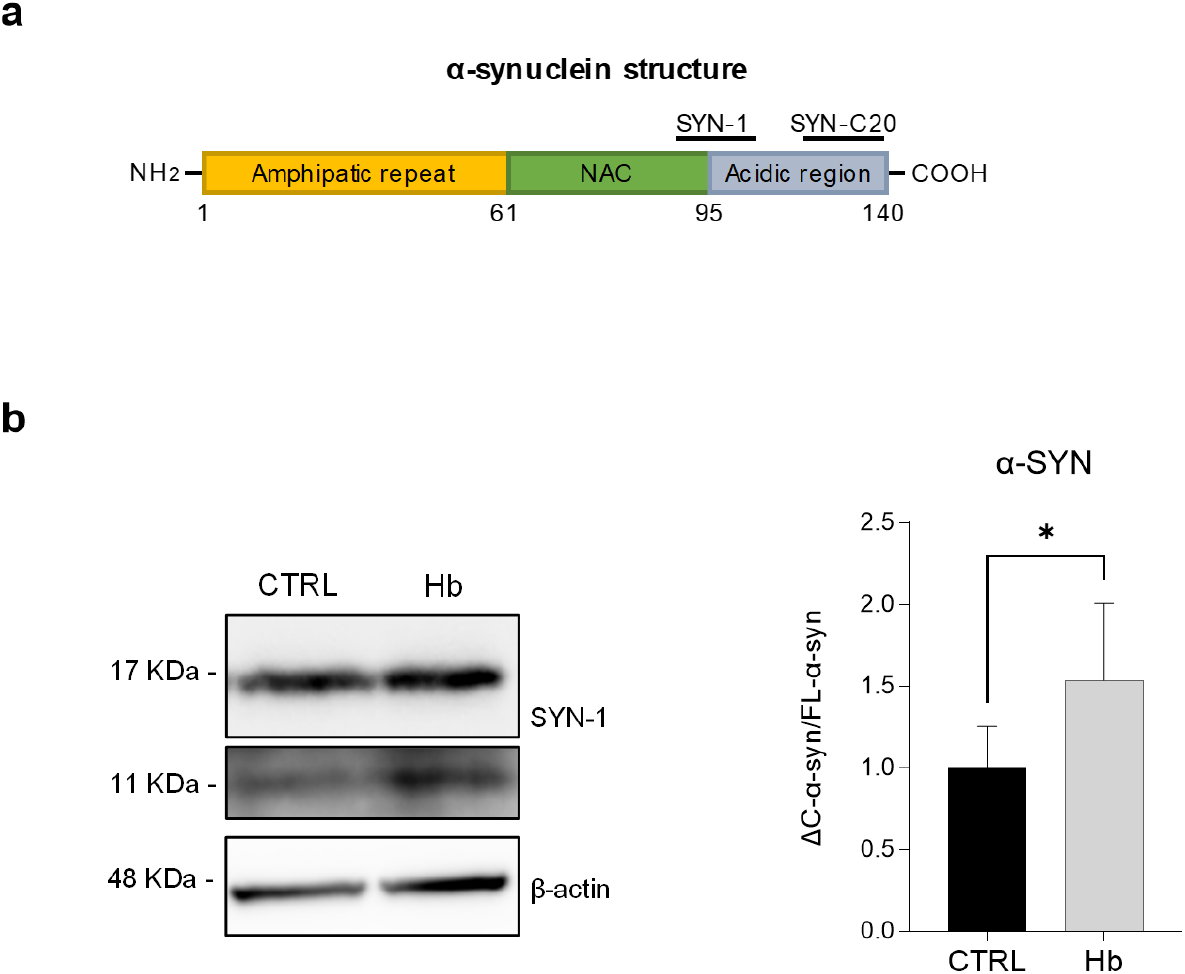
Hb overexpression in SNpc triggers the formation of ΔC-α-syn. Schematic representation of the domains structure of human α-syn protein and epitopes of anti-α-syn antibodies used for the study (**a**). FL-α-syn (17 KDa), ΔC-α-syn (11 KDa) expression in SNpc of Hb and CTRL mice (n=8) at 10 months post-injection of (**b**, *left panel*) was assessed by western blot. Band intensity was quantified (**b**, *right panel*).

C-terminal cleavage is one of the major post-translational modifications (PTM) of α-syn. It could be particularly detrimental as ΔC-α-syn self-assembles into fibrils and increases the aggregation rate and toxicity in both cultured cells [22, 23] and animal models [24–26].

Therefore, in the remaining part of this work we studied the interplay between Hb and α-syn and the formation of α-syn C-terminal fragments *in vitro*.

### Biochemical analysis and structural characterization of α-syn preformed fibrils (PFFs) preparation

To study the role of Hb in α-syn C-terminal truncation, we first prepared PFFs to induce α-syn misfolding cyclic amplification in Hb-overexpressing and control cell lines. Recombinant human α-syn was expressed in *E.coli* BL21(DE3) strain and purified. Fibrillation reactions were monitored by thioflavin T (ThT) fluorescence and PFFs were collected at plateau as long fibrils (Supplementary Figure 2a). Atomic force microscopy (AFM) was performed as previously reported [27] to confirm the formation of PFFs (Supplementary Figure 2b). The presence of high molecular weight α-syn species in PFFs preparations was detected as smear by Western Blot. Both Monomers (Ms) and PFFs preparations contained monomeric and dimeric α-syn. In addition, both preparations showed ΔC-α-syn species (Supplementary Figure 2d).

As a cellular model system, we took advantage of Hb cells, the DA iMN9D mouse cell line stably overexpressing α and β-globins and forming α2β2 tetramer, as previously described [8–10]. CTRLs were iMN9D cells stably transfected with a control vector [8, 9, 10, 28].

Alexa-488 labelled PFFs and Ms were thus supplemented to Hb and CTRL cells for 24 hours. The intracellular accumulation of PFFs, but not of Ms, was confirmed by western blot in both cell lines (Figure 4a). Intracellular fluorescent punctate structures were evident in supplemented cells proving PFFs internalization (Figure 4b). This results was confirmed by a modified IF assay in which cells were incubated with Trypan blue, a known green fluorescence quencher with an affinity for amyloid (*data not shown*) [29]. Importantly, PFFs inclusions were stained with an antibody for pSer129 (Figure 4b), the prominent PTMs involved in α-syn fibrillation [30, 31].

**Figure 4.**
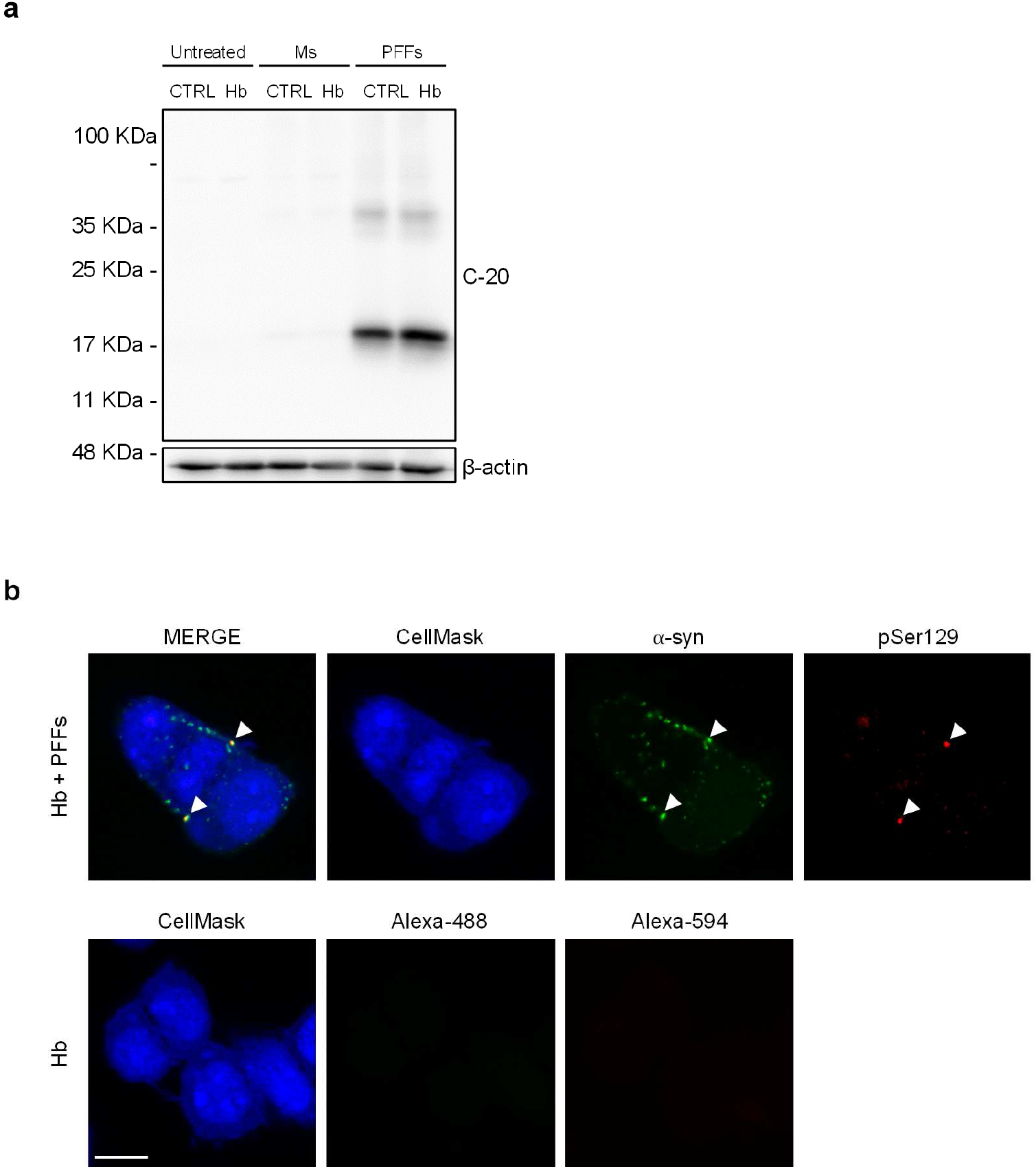
Sonicated α-syn fibrils are internalized by Hb cells and are positively stained by pSer129 antibody. Representative western blot of lysates from Hb and CTRL cells upon supplementation with vehicle, Ms and PFFs for 24 h **(a)**. Representative confocal microscopy images of Hb cells supplemented with Alexa-488 labelled PFFs for 24 h. IF for α-syn phosphorylated at Ser129 (pSer129, Alexa 594, red) is shown. Arrows indicate intracellular inclusions positive to pSer129 antibody. Entire cells were labelled by CellMask. Untransfected cells incubated only with secondary antibody were used to establish autofluorescence levels. Nuclei were stained with DAPI. Scale bar 10 μm.

### Hb triggers the accumulation of a C-terminal truncated form of α-syn in vitro

Hb and CTRL cells were then supplemented with PFFs for 24, 48 and 96 hours and the pattern of α-syn species was studied by western blot analysis using SYN-1 or SYN-C-20 antibodies. In general, the intracellular expression of FL-α-syn decreased over time, whereas ΔC-α-syn species increased (Figure 5a and b). In contrast, the levels of extracellular α-syn species remained stable (Supplementary Figure 3). Notably, ΔC-α-syn was reproducibly more abundant in Hb than CTRL cells at each time point and the levels of ΔC-α-syn normalized to FL-α-syn (ΔC-α-syn/FL-α-syn ratio) were higher in Hb than CTRL cells with a statistically significant difference at each time point (Figure 5a and c). These results recapitulate *in vitro* what observed upon Hb overexpression in mouse SNpc strengthening the significance of the findings.

**Figure 5.**
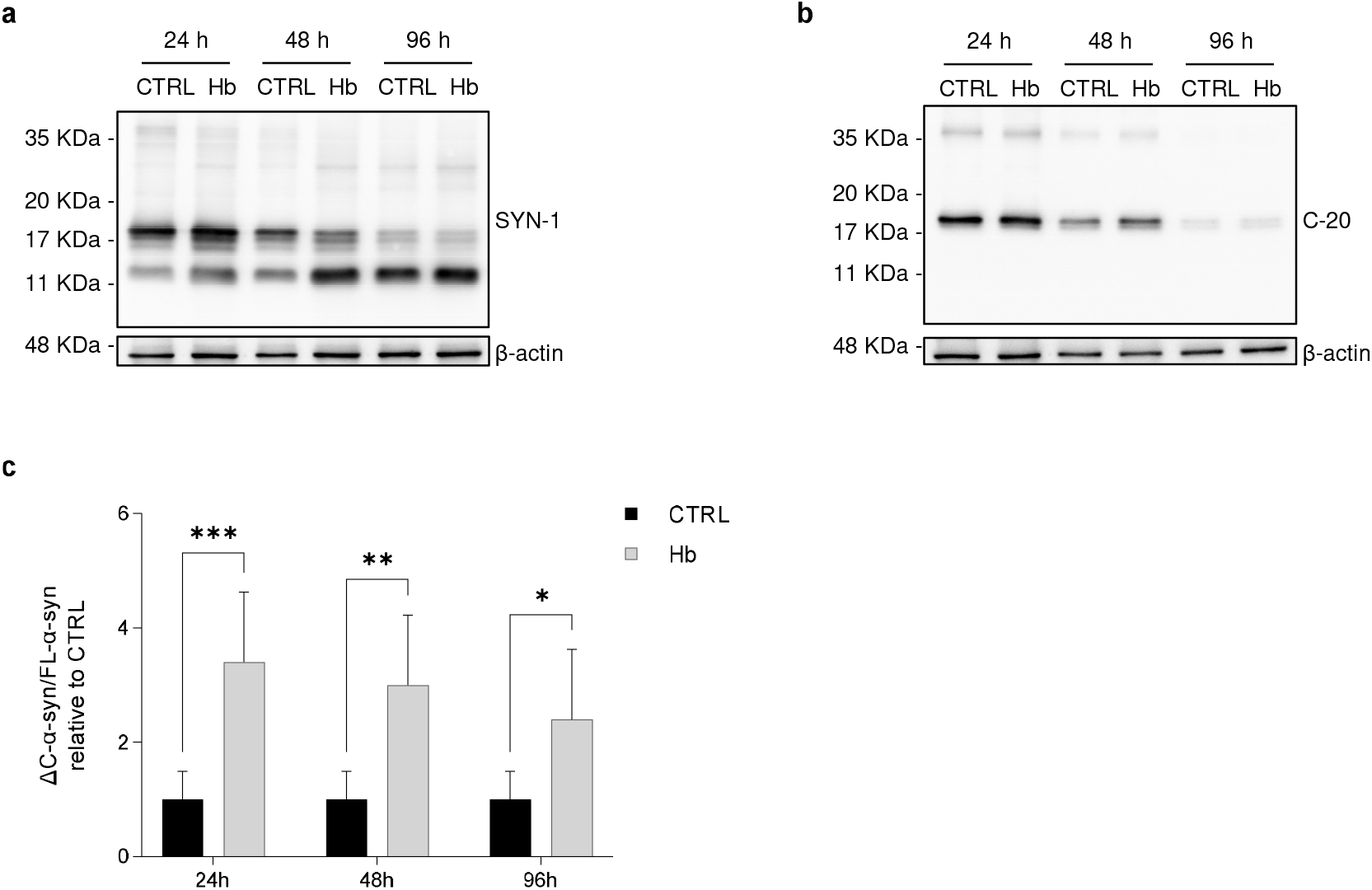
C-terminal truncated α-syn accumulation in the presence of Hb in cell lysates. CTRL and Hb cells were treated with α-syn amyloids. Cell lysates were collected at the indicated time points. Cell lysates were analysed by immunoblotting with SYN-1 (**a)**and C-20 antibodies (**b**). Band intensity corresponding to ΔC-α-syn and FL-α-syn was quantified and the ratio was calculated. Data represent means ± SEM and are representative of six independent experiments. Statistical analysis was performed with one-way Anova. *, p ≤ 0.05; **, p ≤ 0.01; ^***^, p≤ 0.001; ****, p ≤ 0.0001 (**c**).

Given the reported presence of Hb-α-syn complexes in primate brains and red blood cells, we investigated their formation in our experimental settings by incubating Hb and CTRL cell lysates with biotinylated PFFs and by puling-down fibrils through NeutrAvidin resin. WB analysis of fibrils did not reveal the co-precipitation of Hb, a finding that exclude the ability of PFFs to bind Hb *per se* (Supplementary Figure 4).

### Proteases contribution to the accumulation of α-syn C-terminal truncated species

To identify the endogenous protease/s that cleave/s α-syn in our experimental settings, RNA-seq data of Hb cells were interrogated [8] and showed that these cells mainly express Calpain I (Capn I) and Cathepsin D (Ctsd), making them candidates for this activity.

To investigate the role of Capn I, we treated Hb cells with the specific inhibitor Capn inhibitor III (CI-III) and measured ΔC-α-syn/FL-α-syn ratio. Hb cells were pre-treated with 10 μM CI-III and the vehicle (DMSO) for 24 h. Medium was then removed and replaced with a new one containing PFFs and CI-III at the same concentration as used before. Cells were then collected after additional 24 or 48 hours for western blot analysis.

ΔC-α-syn/FL-α-syn ratio decreased in a statistically significant manner upon 48 hours of CI-III treatment, demonstrating that Capn I is involved in α-syn C-terminal truncation in our experimental settings (Figure 6a, b and c). Cells treated with CI-III also showed increased levels of ⍺-spectrin, a well-known Capn I substrate, to confirm Calpain inhibitor activity (Figure 6d).

**Figure 6.**
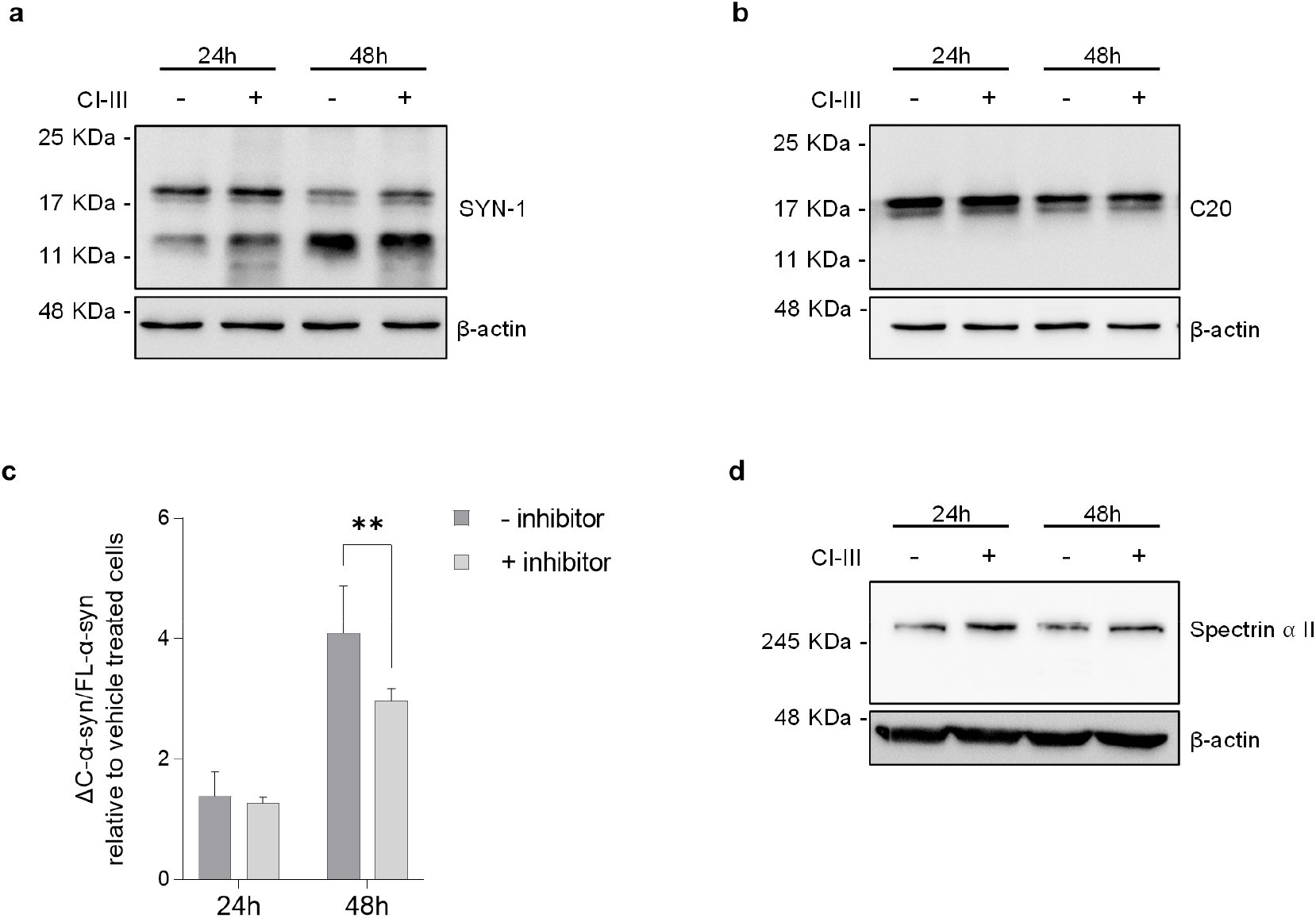
Effect of Calpain I inhibition on α-syn C-terminal truncated species accumulation in Hb cells. Cell lysates of Hb cells treated with DMSO (−) and Calpain inhibitor III (+) were analysed by immunoblotting with SYN-1 (**a**) and C-20 (**b**) antibodies. Band intensity corresponding to ΔC-α-syn and FL-α-syn was quantified and the ratio was calculated. Data represent means ± SEM and are representative of six independent experiments. Statistical analysis was performed with one-way Anova. *, p ≤ 0.05; **, p ≤ 0.01; ^***^, p≤ 0.001; ****, p ≤ 0.0001 (**c**). Cell lysates of Hb cells treated with DMSO (−) and CI-III (+) were analysed by immunoblotting with anti-Spectrin α II antibody (**d**).

The very same experimental settings were then used to assess the role of Ctsd. As shown in Supplementary Figure 5, 100 μM of Pepstatin A, an inhibitor of acid proteases including Ctsd, did not modify the pattern of α-syn truncation. It should be considered that in our experimental conditions Pepstatin A was able to inhibit only 20 or 30% of Ctsd activity respectively at 24 and 48 hours upon PFFs supplementation (Supplementary Figure 5; see Material and Methods). Given that higher Pepstatin A concentration led to cell death, additional experiments are needed to definitively rule out Ctsd as involved in Hb-induced α-syn fragments formation. These results show that the formation of α-syn fragments induced by Hb depends, at least in part, by Calpain activity, a protease previously shown to be involved in α-syn PTM.

## DISCUSSION

Hb is attracting interest in the field of PD for some intriguing observations: i. Hb expression in the mesencephalic DA cell system correlates to cell’s vulnerability in disease [8]; ii. Hb is mainly, but not exclusively, localized in the mitochondria, an organelle crucial for PD pathogenesis, and it influences its activity [8,12]; iii. its association to mitochondria decreases in aging and post-mortem PD brains [13–15]; iv. Hb overexpression increases DA cells’ susceptibility to neurochemical intoxication, a PD cellular model [9]; v. Hb inhibits stress-induced autophagy, a pathway involved in PD [9].

Here, we show that long-term overexpression of Hb in mouse SN leads to a loss of 50% of DA neurons after 9 months from injection. We previously showed that after one month, Hb was forming intracellular aggregates localized to the cytoplasm and to the nucleolus leading to initial impairment of motor learning with no cellular loss [9]. Here, we monitored AAV-injected mice for behavioural deficits along the entire timeframe of the experiment since the onset of behavioural alterations was expected to be slow and progressive. Hb mice presented subtle motor deficits and evident cognitive dysfunctions. Locomotor activity and rotarod test did not evidence overt motor impairments. However, two tests that evaluate different motor skills showed a decrease in abilities in Hb mice. These results are in agreement with what is observed in PD patients and animal models, where motor symptoms appear when most of the dopaminergic neurons are lost [1]. Hb mice showed substantial defects in memory assessment and spatial working memory. This is consistent with the fact that cognitive impairments usually appear in PD patients before the onset of motor symptoms [32]. Similar deficits have been observed in a PD mouse model with bilateral partial 6-hydroxydopamine lesions and loss of SNpc DA neurons of about 60%. These mice showed a mild motor phenotype (e.g. no locomotor activity alteration) and cognitive deficits, as evidenced in NOR test and in other behavioral assays not related to motor functions [33]. It is therefore intriguing that the overexpression of Hb is phenocopying features of a well-accepted neurochemical model of PD.

It remains unclear how Hb is triggering DA cells’ loss. To answer this question, here we provide evidence that Hb overexpression *in vivo* and *in vitro* triggers ΔC-α-syn formation. Pathologic α-syn presents several PTMs. Prominent features of the protein in the inclusion bodies are the phosphorylation at serine 129 (pSer-129) and C-terminal truncations [30, 34]. ΔC-α-syn species are known to increase the pathological aggregation into LB inclusions because of the propensity to aggregate themselves, to promote the aggregation of FL-α-syn and to trigger toxicity. Such a prion-like seeding mechanism of ΔC-α-syn has been observed both *in vitro* and *in vivo* [35–38].

In post-mortem PD brains, LB inclusions in SNpc neurons comprise a 20% of amyloidogenic ΔC-α-syn [1, [39, 40]. Since α-syn fragments accumulate in the vermiform appendix, the fact that patients with appendectomy are less likely to develop PD or present a delayed disease onset, is considered in support of the notion that α-syn carboxy truncations might be relevant in disease pathogenesis [41]. The control of the proteolytic cleavage of α-syn is therefore a crucial event and a viable target for therapeutics.

Endogenous ΔC-α-syn content in SNpc of Hb mice showed an increase of about 50% compared to CTRL mice. These results were recapitulated *in vitro* where iMN9D cells overexpressing α and β chains of Hb were challenged with PFFs [4, 7]. PFFs were internalized by Hb iMN9D cells and promoted aggregation of intracellular pSer129-modified α-syn. ΔC-α-syn/FL-α-syn ratio increased significantly in iMN9D Hb cells as compared to control and provided further evidence that Hb expression correlates with the C-terminal truncation of α-syn. The repertoire of candidate proteases involved in the formation of ΔC-α-syn comprises neurosin, Capn I, Ctsd and matrix metalloproteinase-3 [37]. Our data show that Hb leads to activation of calpain I, since ΔC-α-syn is significantly inhibited by CI-III. Interestingly, calpain inhibitors were shown to exert disease-modifying activity and reduction of α-synuclein deposition in transgenic models of PD [42]. Additional experiments should address the involvement of ctsd since its inhibitor, Pepstatin A, is highly toxic in our experimental setting. It remains unclear how Hb is linked to calpain activation. Given the observation that Hb-α-syn complexes can accumulate in the cytoplasm of striatal and SN cells in monkeys, in human post-mortem brains and in red blood cells [13, 14], we investigated whether Hb and α-syn were forming complexes in our experimental conditions. No interactions were observed leaving the mechanistic link between Hb, calpain activation and α-syn cleavage unclear. Importantly, the formation of endogenous ΔC-α-syn fragments could be a crucial step in the chain of events from Hb expression to DA cell’s loss *in vivo*. In support of this model, loss of DA neurons has been observed in mice overexpressing ΔC-α-syn [43, 44]. They also showed deficits in locomotion and in cortical-hippocampal memory test [24]. Moreover, passive immunization against ΔC-α-syn ameliorated neurodegeneration and neuroinflammation, reduced the accumulation of ΔC-α-syn and improved motor and memory deficits in a mouse model of PD [45]. It will be interesting to investigate whether the amount of endogenous ΔC-α-syn in Hb mice is sufficient to promote neurodegeneration or it requires a second pathological event triggered by Hb.

Our results may be of interest beyond PD. The presence of inclusion bodies rich in fibrillar FL-α-syn and ΔC-α-syn fragments is a common characteristic of Synucleinopathies, a group of neurodegenerative diseases that include Lewy Body Dementia (LBD) and Multiple System Atrophy (MSA). In MSA, the typical α-syn-containing glial cytoplasmic inclusions are restricted to oligodendrocytes [46]. Interestingly, Hb is expressed in mouse and human oligodendrocytes *in vitro* and *in vivo* [8] and MSA mouse models and post-mortem brains show an increase of Hb expression of 2.5 to 3 fold [47, 48].

These results suggest the need for further studies on the role of Hb in these neurodegenerative diseases.

## CONCLUSION

Our study provides new evidence that implicates Hb in PD. Given the effects of Hb overexpression on DA cells’ viability and α-syn metabolism, an analysis of the correlation between genetic variation of Hb genes and Hb levels in the brain is needed to potentially associate its expression to the onset of synucleinopathies, including PD.

## ABBREVIATIONS

α-syn: alpha-synuclein;
Ms: monomers;
PTMs: post-translational modifications;
pSer-129: phosphorylated serine 129;
ΔC-α-syn: C-terminal truncated αsyn;
FL-α-syn: full-length α-syn;
PD: Parkinson’s disease;
LBD: Lewy Body Dementia;
MSA: Multiple System Atrophy;
SNpc: *Subtantia nigra pars compacta*;
GCIs: Glial cytoplasmic inclusions;
DA: dopaminergic;
Hb: hemoglobin;
nHb: neuronal Hb;
MPP^+^: 1-methyl-4-phenylpyridinium;
PFFs: pre-formed fibrils;
AFM: Atomic force microscopy;
NHP: non-human primates;
Capn I: Calpain I;
Ctsd: cathepsin D;
Cas1: caspase 1;
TH: tyrosine hydroxylase;
NOR: novel object recognition;

## ACKNOWLEDGEMENTS

We are indebted to all the members of the SG laboratory for thought-provoking discussions. We are grateful to SISSA, IIT technical and administrative staff, especially to Micaela Grandolfo, Omar Peruzzo and Eva Ferri, and Università del Piemonte Orientale (UPO).

## AVAILABILITY OF DATA AND MATERIALS

The manuscript has data included as electronic Additional information.

## ETHICS APPROVAL

All animal experiments were performed in accordance with European guidelines for animal care and following Italian Board Health permissions (D.Lgs. 26/2014, 4 March 2014).

## AUTHORS’ CONTRIBUTIONS

CS designed, carried and analysed the *in vitro* experiments, wrote the manuscript; CB performed and analysed the *ex vivo* experiments, wrote the manuscript; ED carried out and analysed α-syn recombinant production and fibrillation; MC, ClS and FP analysed the data and discussed experimental results; NJ, PP and PF discussed experimental results; PP performed and PF supervised AFM experiments; GL provide the fibrils, analysed the data and discussed experimental results; S.E. conceived the project and carried out the *in vivo* experiments, analysed the data and composed the manuscript; SG conceived the project, designed the experiments, supervised the study, and wrote the manuscript. All authors contributed to this work, read the manuscript and agreed to its contents.

## AUTHOR DETAILS

*Affiliations:*

^1^Area of Neuroscience, Scuola Internazionale Superiore di Studi Avanzati (SISSA), Trieste, Italy.

^2^Central RNA Laboratory, Istituto Italiano di Tecnologia (IIT), Genova, Italy.

^3^ELETTRA Synchrotron Light Source, Trieste, Italy.

^4^Department of Health Sciences and Research Center on Autoimmune and Allergic Diseases (CAAD), University of Piemonte Orientale (UPO), Novara, Italy.

*Corresponding authors:*

Correspondence to Stefano Gustincich or Stefano Espinoza.

