## Additional Information for "Neuronal hemoglobin induces loss of dopaminergic neurons in mouse Substantia nigra, cognitive deficits and cleavage of endogenous α-synuclein"

### Supplementary Figure 1

a

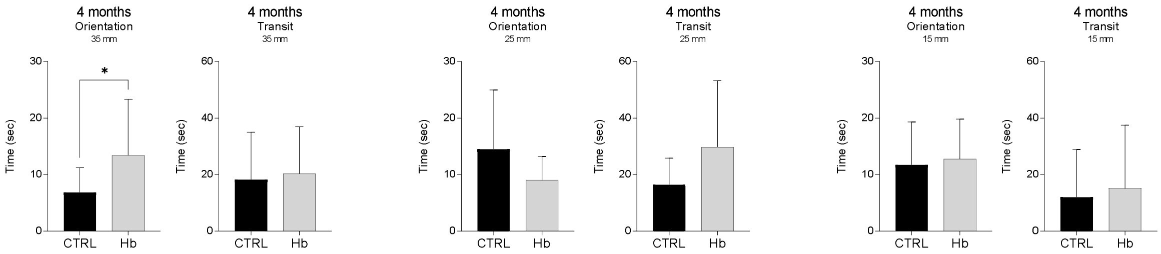


b

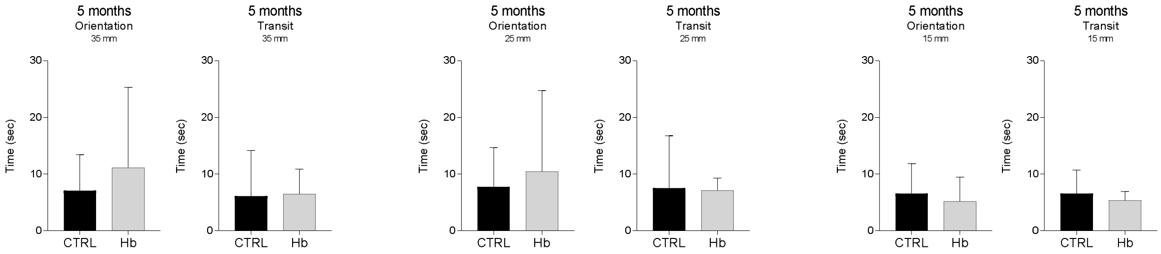


c

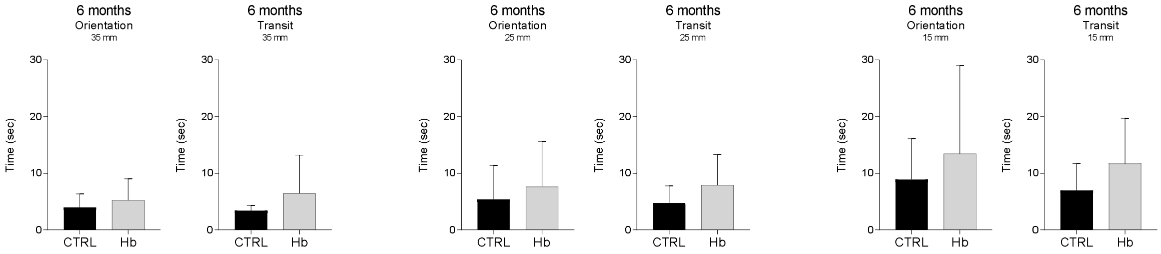


d

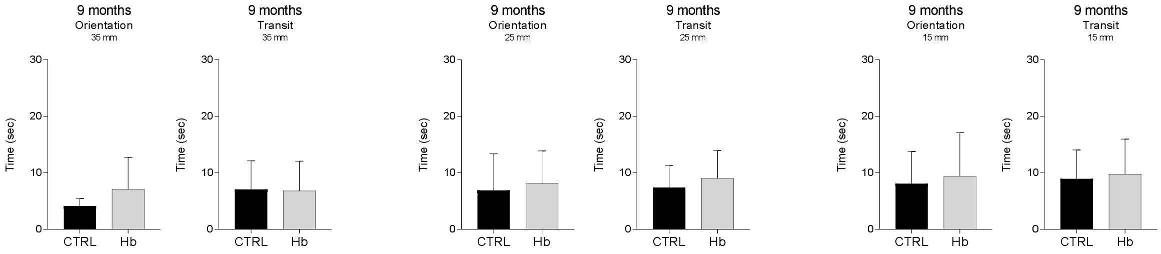


**Static rods test on CTRL and Hb mice.** CTRL (n=15) and Hb (n=15) mice were assessed in static rods test measuring two parameters, transit time and orientation time (seconds) and different time points. Different diameters of the rods were used, 35, 25 and 15mm. The time points evaluated in this test were 4 (**a**), 5 (**b**), 6 (**c**) and 9 months (**d**) after injection. Data represent means ± SEM. Statistical analysis was performed with unpaired t test with Welch's correction. *, $p \leq 0.05$.

### Supplementary Figure 2

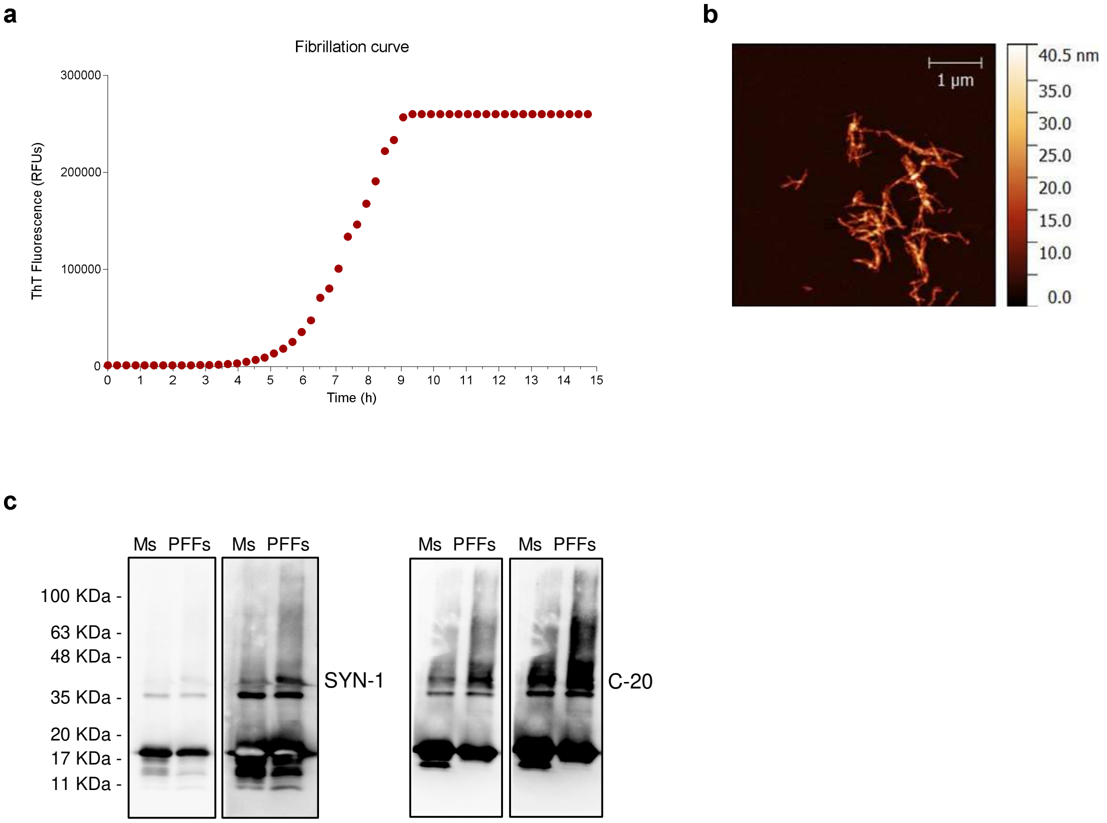


**Biochemical analysis and structural characterization of α-syn preparations.** Fibrillation curve of recombinant human α-syn protein analysed using ThT fluorescence assay. Mean of three wells is represented (**a**). AFM analysis of fibrillary human α-syn aggregates after 5 min of sonication (**b**). Ms and PFFs preparations were analysed by western blot with SYN-1 and C20 antibodies. The same membrane was exposed for a longer period in order to detect high molecular weight species (**c**).

**Supplementary Figure 3**

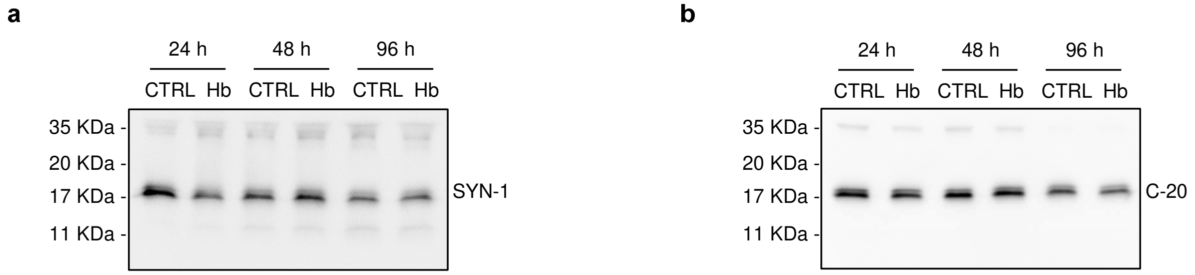


**Accumulation of α-syn in cell media.** CTRL and Hb cells were treated with α-syn amyloids and cell medium were collected at the indicated time points and analysed by immunoblotting with SYN-1 (**a**) and C-20 (**b**) antibodies.

**Supplementary Figure 4**

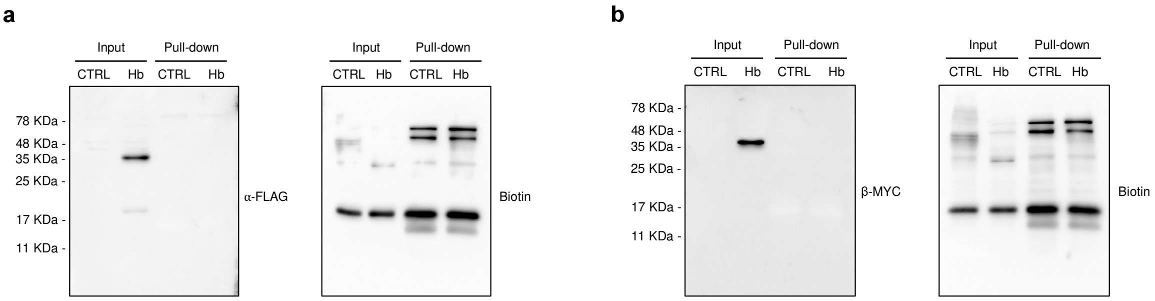


**α-syn and Hb don't form complexes.** Pull down of biotinylated PFFs in cell lysates from both CTRL and Hb cells were revealed by immunoblot with anti-FLAG (**a**) and anti MYC (**b**) antibodies. Samples were also revealed with anti-biotin antibody as control.

**Supplementary Figure 5**

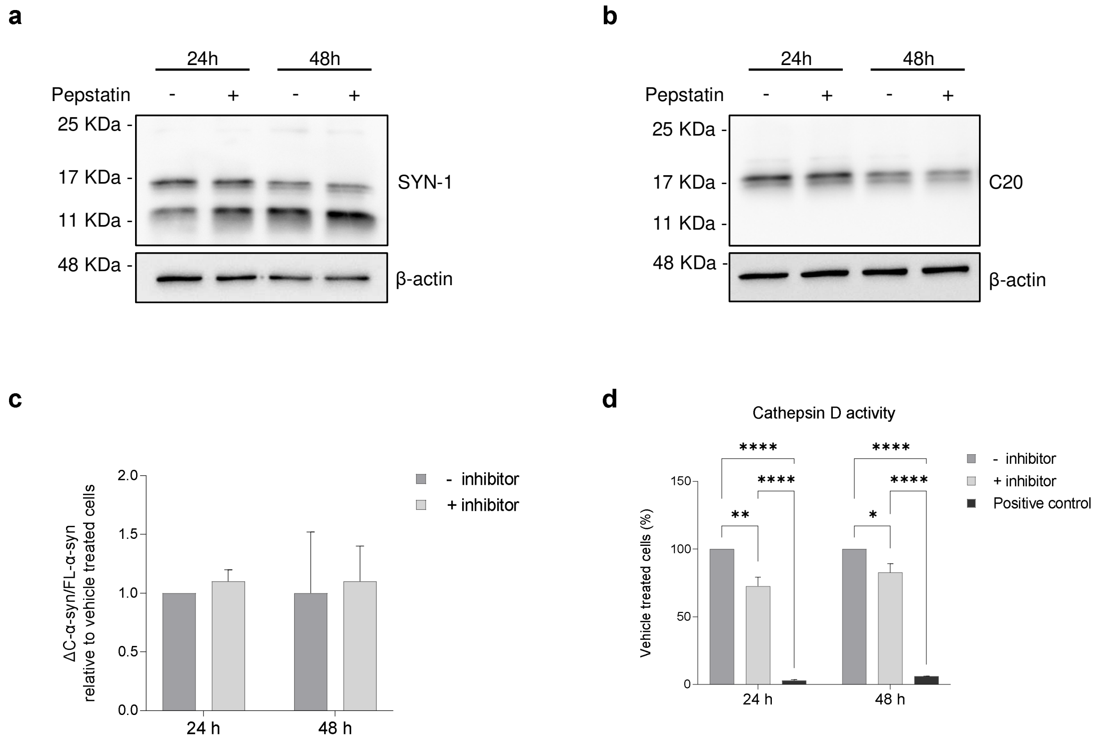


**Effect of Ctsd inhibition on α-syn C-terminal truncated species accumulation in Hb cells.** Cell lysates of Hb cells treated with DMSO (-) and Pepstatin A (+) were analysed by immunoblotting with SYN-1 (**a**) and C-20 (**b**) antibodies. Band intensities corresponding to ΔC-α-syn and FL-α-syn were quantified and the ratio was calculated. Data represent means ± SEM and are representative of five independent experiments. (**c**). Cell lysates of Hb cells treated with DMSO (-) and Pepstatin A (+) were analysed by Ctsd activity assay. Data represent means ± SEM and are representative of two independent experiments, each performed in three replicas and are expressed as percentage of vehicle treated cells (**d**). Statistical analysis was performed with one-way Anova. *, $p \leq 0.05$; **, $p \leq 0.01$; ***, $p \leq 0.001$; ****, $p \leq 0.0001$.
